# Single-cell Transcriptomic Analysis of Systemic Autoinflammatory Diseases with Anti-TNFα Therapy

**DOI:** 10.1101/2022.11.29.518333

**Authors:** Yichao Hua, Na Wu, Junke Miao, Min Shen

**Author notes:** **Correspondence:** Min Shen, Yichao Hua.

## Abstract

Systemic autoinflammatory diseases (SAIDs) are a group of rare diseases characterized by recurrent or continuous inflammation, typically accompanied by genetic variants. Good responses to anti-TNFα therapy were observed in SAIDs patients. However, the mechanisms underlying the disease flare and the response to TNFα blocking therapy have not been fully elucidated. Here, we use single-cell RNA sequencing technology to describe the transcriptomic profile of PBMCs and PMNs in SAIDs patients both before and after anti-TNFα treatment. We found that interferon responses are involved in the active phase of the disease. After anti-TNFα therapy, we observed remission from clinical symptoms while unexpected elevation of TNFα and IL-1, indicating that these inflammatory cytokines are not positively correlated with disease activity. Trajectory analysis showed that inhibition of macrophage differentiation, rather than reduction of the inflammatory cytokines, as the potential mechanism of anti-TNFα treatment response in SAIDs.

## 1 Introduction

Systemic autoinflammatory diseases (SAIDs) are a group of rare diseases caused by defects or dysregulation of the innate immune system, characterized by recurrent or continuous inflammation (e.g. fever, serosal, synovial, or cutaneous inflammation) and by the lack of a primary pathogenic role of the adaptive immune system (autoreactive T-cells or autoantibodies production) (Ben-Chetrit et al., 2018; Centola et al., 1998; Manthiram et al., 2017; Nigrovic et al., 2020). This group of diseases includes more than 40 monogenic, as well as several polygenic diseases (Doria et al., 2012; de Jesus et al., 2015). Classic monogenic SAIDs include familial Mediterranean fever (FMF, caused by *MEFV* gene variants), TNF receptor-associated periodic fever syndrome (TRAPS, by autosomal dominant variants in the *TNFRSF1A* gene), cryopyrin-associated periodic syndromes (CAPS)/*NLRP3*-associated autoinflammatory disease (*NLRP3-AID*, by gain-of-function variants in the *NLRP3* gene), and hyper-IgD syndrome (HIDS)/mevalonate kinase deficiency (MKD, by autosomal recessive variants in the *MVK* gene) (de Jesus et al., 2015). Patients with high disease activity can present with recurrent attacks of fever and other inflammatory symptoms, and a successful treatment can reduce disease activity, as manifested by reduced frequency or severity of disease flare or no flare. Regarding to pathogenesis, these classic SAIDs are classified as IL-1 mediated SAIDs, as the variants can directly or indirectly promote the assembly of inflammasomes that proteolytically process the inactive pro-IL-1β into active IL-1β. This can occur by increased intracellular sensor/pattern-recognition receptors (PRR) function (e.g. CAPS, FMF), generation of intracellular stress (e.g. TRAPS, HIDS), or loss of a negative regulator (e.g. Deficiency of IL-1 receptor antagonist, DIRA) (de Jesus et al., 2015). A lot of studies focus on the pathogenic role of the inflammasome – IL-1 axis in these SAIDs, and many patients have good responses to IL-1-blocking agents (e.g. anakinra, canakinumab, and rilonacept), although some patients with FMF, TRAPS or HIDS are less responsive to IL-1 inhibition (Cantarini et al., 2015; Dinarello and van der Meer, 2013; Hoffman et al., 2008, 2012; Jesus and Goldbach-Mansky, 2014; Kuemmerle-Deschner et al., 2011a, 2011b; Schett et al., 2016; Szekanecz et al., 2021). Little is known about other pathways in the pathogenesis of SAIDs, and the treatment response to other biological agents, e.g. TNFα or IL-6 blocking agents. Some studies have reported increased NFκB activation and defective autophagy as potential pathogenic mechanisms (Bachetti et al., 2013; Hua et al., 2018; Nedjai et al., 2008). In our previous study, we observed a good response to TNFα inhibitors in CAPS patients (Wu et al., 2021). Whether there are other pathways involved in the pathogenesis of SAIDs, and what the underlying mechanisms are in response to TNFα blocking therapy, remain to be elucidated.

Single-cell RNA sequencing (scRNAseq) is a powerful technique to detect transcriptomics of single cells, which can further reveal cellular heterogeneity, gene regulatory networks, cell-cell interactions, and differentiation trajectories in tissues. This technique has been widely used in oncology, microbiology, neurology, and immunology (Tang et al., 2019). Yet, it has never been applied to SAIDs research. In this study, we are the first to apply this technique to classic SAIDs, aiming to identify the potential mechanisms of disease flare other than the IL-1 pathway and the response to anti-TNFα therapy, and to provide the first transcriptomic resource of SAIDs spectrum diseases.

## 2 Results

### 2.1 Single Cell Profiling of PBMCs and PMNs in SAID Patients

This study enrolled 2 patients with classic SAIDs. One patient was a 20-year-old Chinese woman diagnosed as CAPS, with *NLRP3* D303G variation, and another patient was a 16-year-old Chinese man diagnosed as TRAPS, with *TNFRSF1A* V202D variation. Detailed information can be found in **Supplemental Data S1**. Both patients had high disease activity during the first visit (pre-treatment). The CAPS patient presented urticarial rash, arthritis (interphalangeal joint and knees), lymphadenopathy, hearing loss, and vision loss, while the TRAPS patient was in the intermittent phase of disease episodes and did not show significant symptoms. Both patients received Etanercept treatment, and during the 3-month follow-up visit (on-treatment), both patients reported significant remission of clinical symptoms and no disease relapse.

We extracted peripheral blood mononuclear cells (PBMCs) and polymorphonuclear neutrophils (PMNs) from pre-treatment (CAPS_NT, TRAPS_NT) and on-treatment (CAPS_TNFi, TRAPS_TNFi) blood samples of both patients, as well as the blood of one healthy donor (HD). Then the cells were subjected to 3’-scRNAseq (10X Genomics) to study the transcriptomic profiles (**Figure 1A**). As the average detected genes (Ngene) in different cell types varied dramatically (e.g. neutrophils and platelets had very low Ngene, while monocytes and plasma cells had higher Ngene, **Figure 1F**), we used different quality filtering criteria for each cluster (Star Methods) and acquired 30,377 cells for the downstream analysis. To identify the common clusters of all 5 samples, we used the Harmony algorithm (Korsunsky et al., 2019) to remove the batch effect, and identified 17 clusters by unsupervised clustering (**Figure 1B, 1C**), including two B-cell clusters (memory, naïve), plasma cells, five T-cell clusters (memory CD4, naïve CD4, memory CD8, naïve CD8, effector CD8), two NK-cell subtypes (CD16^+^ and CD16^-^), two monocyte subtypes (CD14^+^, CD16^+^), DCs, pDCs, neutrophils, platelets, and progenitors (**Figure 1B**). Representative markers of each cell type were shown in **Figure 1D**. T-cells, CD14 monocytes, CD16 NK-cells, and neutrophils were more frequent cell types in the blood (**Figure 1E**).

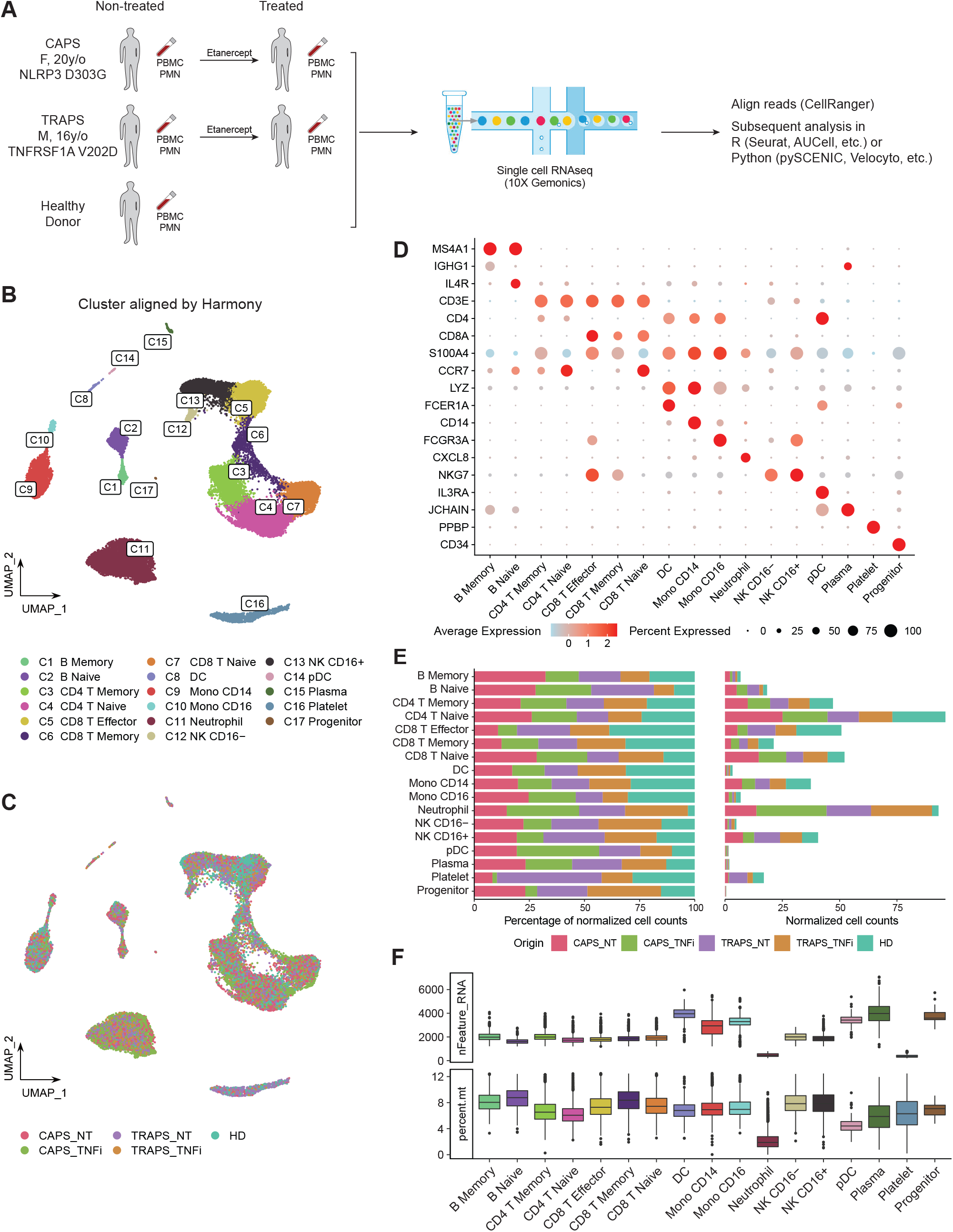

### 2.2 Gene Expression Profile Alters Significantly after Anti-TNFα Therapy in CAPS Patient

After identifying common clusters, we investigated the differences among these samples in each cell type. We plotted unaligned UMAP based on PCA without Harmony integration and it intuitively showed marked changes in the gene expression profile of CAPS after treatment, compared to other samples (**Figure 2A**). This may suggest that the transcriptome is more associated with disease attack rather than disease activity, as both patients had high activity but only the CAPS patient was at disease flare. To make the comparison more quantitative, we calculated the Pearson Correlation of average gene expression of each cluster between groups (**Figure 2B**). Note that low Ngene and cell counts can usually generate lower Pearson Correlation value due to a higher variance, like neutrophils, platelets, DCs/pDCs and CD16^-^ NK cells. In general, CAPS_TNFi shows a lower correlation value compared to other samples, which confirmed our observation in **Figure 2A**. B cells, effector and memory CD8 T cells, NK cells and monocytes had relatively higher variance than naïve or CD4 T cells, indicating that these can be the main cell types that respond to anti-TNFα therapy in the CAPS patient.

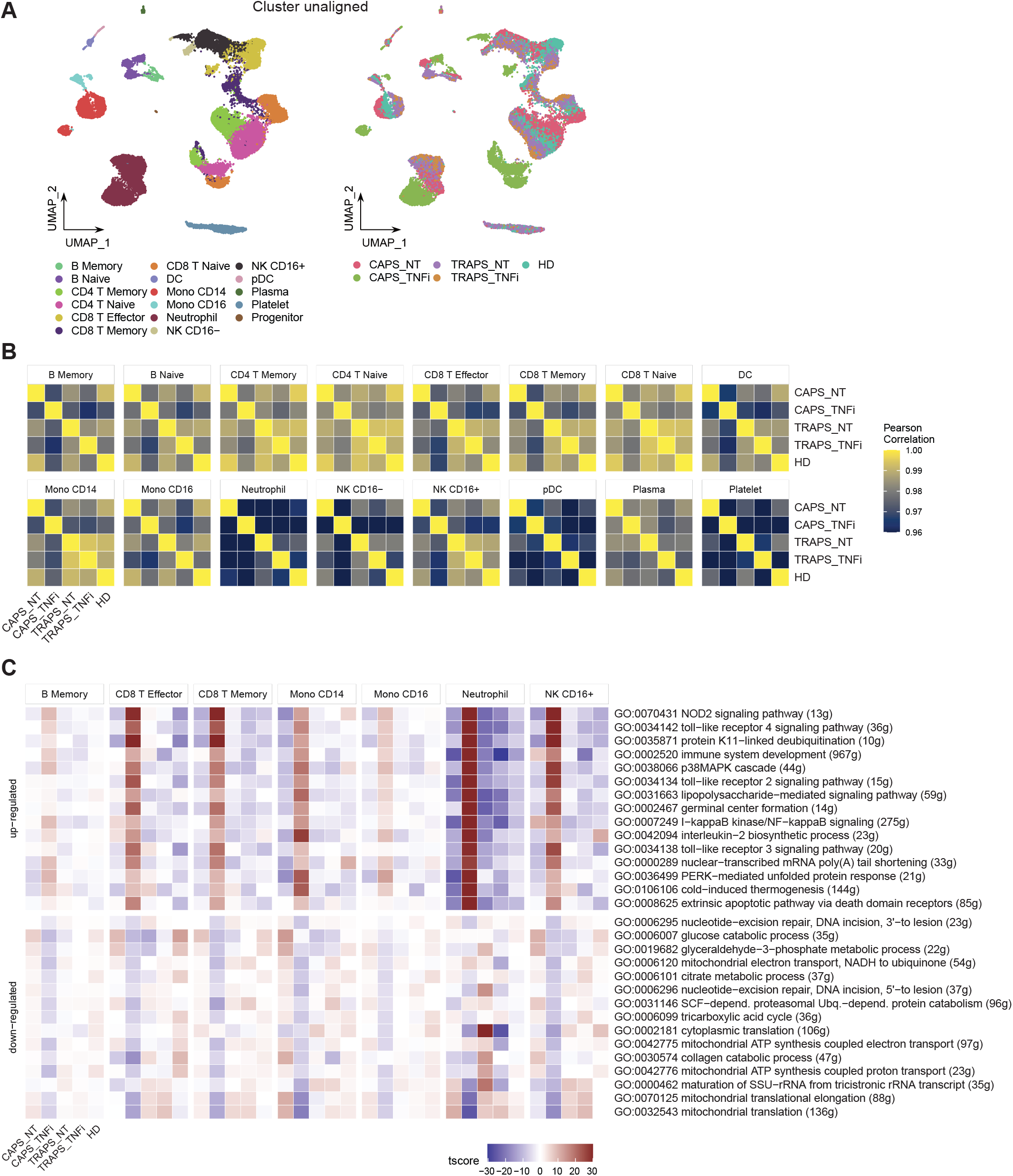

To further look into which pathways contribute to the major changes in CAPS_TNFi, we performed Gene Set Enrichment Analysis (GSEA) based on Gene Ontology (GO) database, and identified common top 10 up- and down-regulated pathways in some representative cell types (**Figure 2C**). Surprisingly, many up-regulated pathways after anti-TNFα therapy in CAPS were related to stress response and innate immune response-activating signal transduction, including pattern recognition receptor (PRR) signaling pathways like toll-like receptor (TLR) and NOD2 signaling pathways. Meanwhile, some metabolic pathways related to aerobic respiration were down-regulated (**Figure 2C**). To confirm this, we performed SCENIC analysis (Aibar et al., 2017) which inferred the gene regulatory network and predicted regulon (transcription factor, TF) activities, and found that indeed many inflammatory-related TFs are activated in CAPS_TNFi, including canonical NFκB pathway TFs (NFKB1, REL) and AP-1 complex (FOS, JUN) (Zenz et al., 2008) (**Figures 3A, B**). We reasoned that the upregulation of inflammatory pathways could be a compensatory adjustment to anti-TNFα therapy, although this requires further validation.

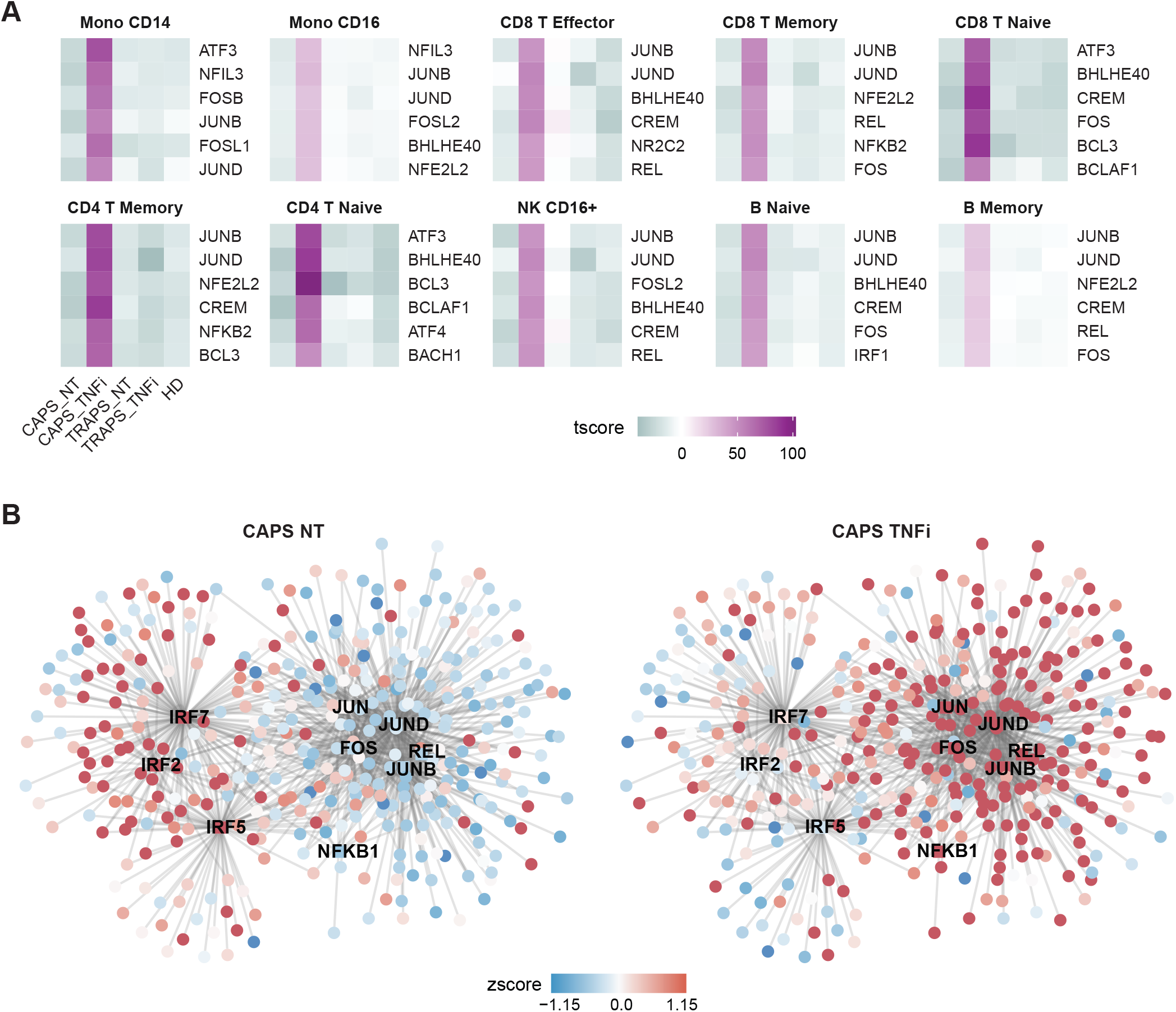

### 2.3 Disease Flare Is Unnecessary to Be Positively Correlated with Serum Inflammatory Cytokines

One important clinical question is which markers are elevated during disease attack. Normally, we evaluate clinical manifestations and laboratory tests including white blood cell (WBC) count, erythrocyte sedimentation rate (ESR), and C-reactive protein (CRP), etc (Singh, 2014). Some inflammatory cytokines, like TNFα, IL-1β and IL-6 are considered to be the cause of systemic inflammation, although IL-6 has both pro- and anti-inflammatory functions (Kany et al., 2019). Thus, we asked how these inflammatory cytokines changed after anti-TNFα therapy. First, we checked which cell types were the main source of each cytokine (**Figure 4A**), and found that TNFα was mainly secreted by CD16^+^ monocytes, IL-1 β mainly by monocytes, DCs and neutrophils, and IL-6 by a small subset of B cells. Next, we checked the expression level of these cytokines in relevant clusters. We noticed that in general they were even more highly expressed after anti-TNFα treatment (**Figure 4B**). This appears counterintuitive, while similar to what we found in **Figure 2C**. Thus, we asked if the increase of inflammatory cytokines after anti-TNFα treatment occurred commonly, by chance, or was just a technical effect of scRNAseq. Therefore, we tried to validate this using other methods. As some patients were tested for serum TNFα and IL-6 during their visit to the hospital, we checked if these cytokines were increased after anti-TNFα treatment at the protein level. We reviewed our SAIDs cohort from 2015 (Hua et al., 2019), until Jan 2021, and among total of 228 patients, 34 were received anti-TNFα therapy (either Etanercept, Infliximab, Golimumab or Adalimumab), and 8 patients were tested for TNFα and IL-6 both before and after anti-TNFα therapy. We found that serum TNFα levels were increased while IL-6 decreased after the treatment (**Figure 4C**). Although it is still not clear what role IL-6 plays during the treatment, we confirmed that TNFα was increased after anti-TNFα therapy, by both scRNAseq and serum protein test. Taken together, the inflammatory cytokines, at least TNFα and IL-1 β can be up-regulated after anti-TNFα therapy and are not necessarily positively correlated with disease flare.

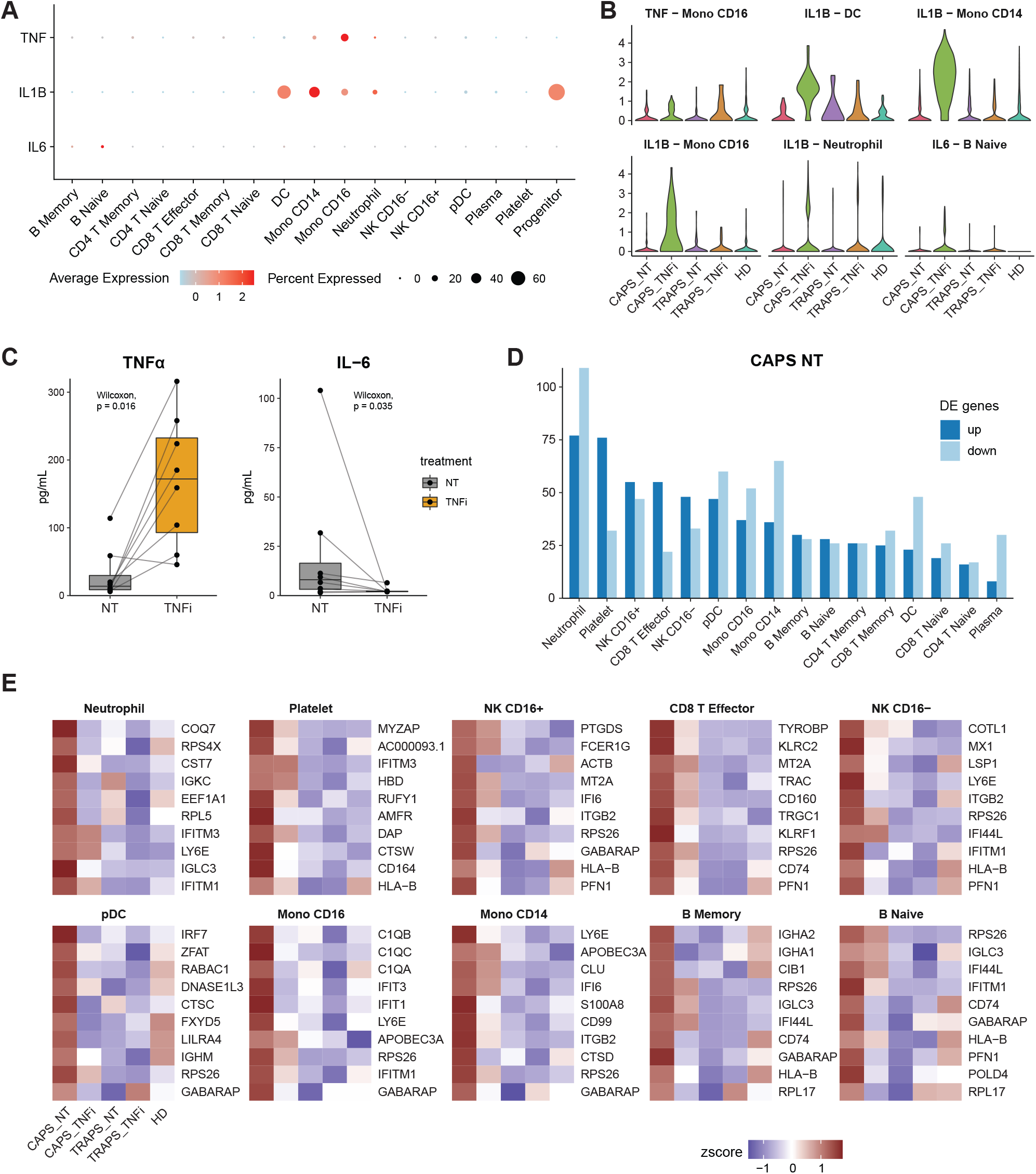

Next, we asked what changes reflect the disease attack, if not TNFα or IL-1β. As we know that among the 5 samples, the CAPS_NT sample was taken during disease attack while other samples were not. Therefore, we checked the differentially expressed genes (DEGs) of CAPS_NT in each cell type (**Figure 4D, E**). We noted that some interferon-induced genes, including IFI, IFIT and IFITM families, which are involved in broad-spectrum antiviral functions (Diamond and Farzan, 2013) were upregulated in CAPS_NT, remarkably in the CD16^+^ monocyte subtype (**Figure 4E**). Consistently, by SCENIC analysis we also observed that several interferon regulatory factors, e.g. IRF2, 5 and 7 were activated in the CAPS_NT sample in CD16 monocytes (**Figure 3B**). These data indicated that the interferon response might play a role in the disease flare.

We further investigated whether the increased interferon response in CAPS_NT was caused by elevated stimuli levels (IFN-α, IFN-γ), or enhanced sensitivity in these monocytes. First, we checked IFN-α and IFN-γ at both RNA and protein levels. In the single-cell RNA-seq data, we did not see increased IFN expression in non-treated samples (**Figure S1A**). At the protein level, we acquired several serum samples of non-treated SAID patients (CAPS = 4, TRAPS = 1) from the biobank in our hospital as well as 3 healthy donors, and tested IFN-α and IFN-γ levels by ELISA. Consistent with scRNAseq data, there was no increased serum IFN in non-treated SAID samples (**Figure S1B**). We then checked the IFN signaling pathways in CD16 monocytes. We used the IFN signaling gene lists from the Reactome database (Gillespie et al., 2022) and calculated the enrichment score using the AUCell package (Aibar et al., 2017). We found that indeed there was higher activation of IFN signaling pathways in CAPS_NT (**Figure S1C**). These data supported that the increased interferon response in CAPS_NT was probably caused by enhanced sensitivity to the IFN stimuli in CD16 monocytes, rather than elevated ligand levels.

### 2.4 Inhibition of Macrophage Differentiation Is a Potential Mechanism of Anti-TNFα Therapy

From the previous analysis, we also noticed that complement components (C1Qs) were more upregulated in pre-treatment than on-treatment samples in CD16^+^ monocytes (**Figure 4E**), which is one of the key features of macrophages (Hartung and Hadding, 1983). Interestingly, CD16^+^ monocytes indeed resemble macrophages in many aspects, as they have low expression of classic monocyte markers like CD14 and CCR2, and highly express CX3CR1 which explains why they migrate and adhere more than CD16^-^ monocytes to fractalkine-secreting endothelium, and specialize in complement and FcR-mediated phagocytosis and anti-viral responses (Davies et al., 2013; Italiani and Boraschi, 2014; Kapellos et al., 2019). The lower expression in complement after TNFα blockade gave us a hint that the treatment might change the differentiation state of CD16^+^ monocytes. To validate this hypothesis, we performed pseudo-time trajectory analysis within monocyte clusters using Palantir and scVelo/Velocyto (Bergen et al., 2020; La Manno et al., 2018; Setty et al., 2019) which could predict the direction of differentiation and “RNA velocity” — the time derivative of the gene expression state — based on the relative abundance of nascent (unspliced) and mature (spliced) mRNA (**Figure 5A, 5B**). We found that before treatment, the CD16^+^ monocytes were highly dynamic and differentiating into the terminal state, while after TNFα blocking, the differentiation was blocked, depicted by the arrows with shorter lengths than untreated (**Figure 5B**). Indeed, TNFα is known to be involved in macrophage differentiation (Witsell and Schook, 1992). This analysis shows that inhibition of macrophage differentiation, rather than reducing the overall level of inflammatory cytokines, could be a potential mechanism of anti-TNFα treatment response in SAIDs.

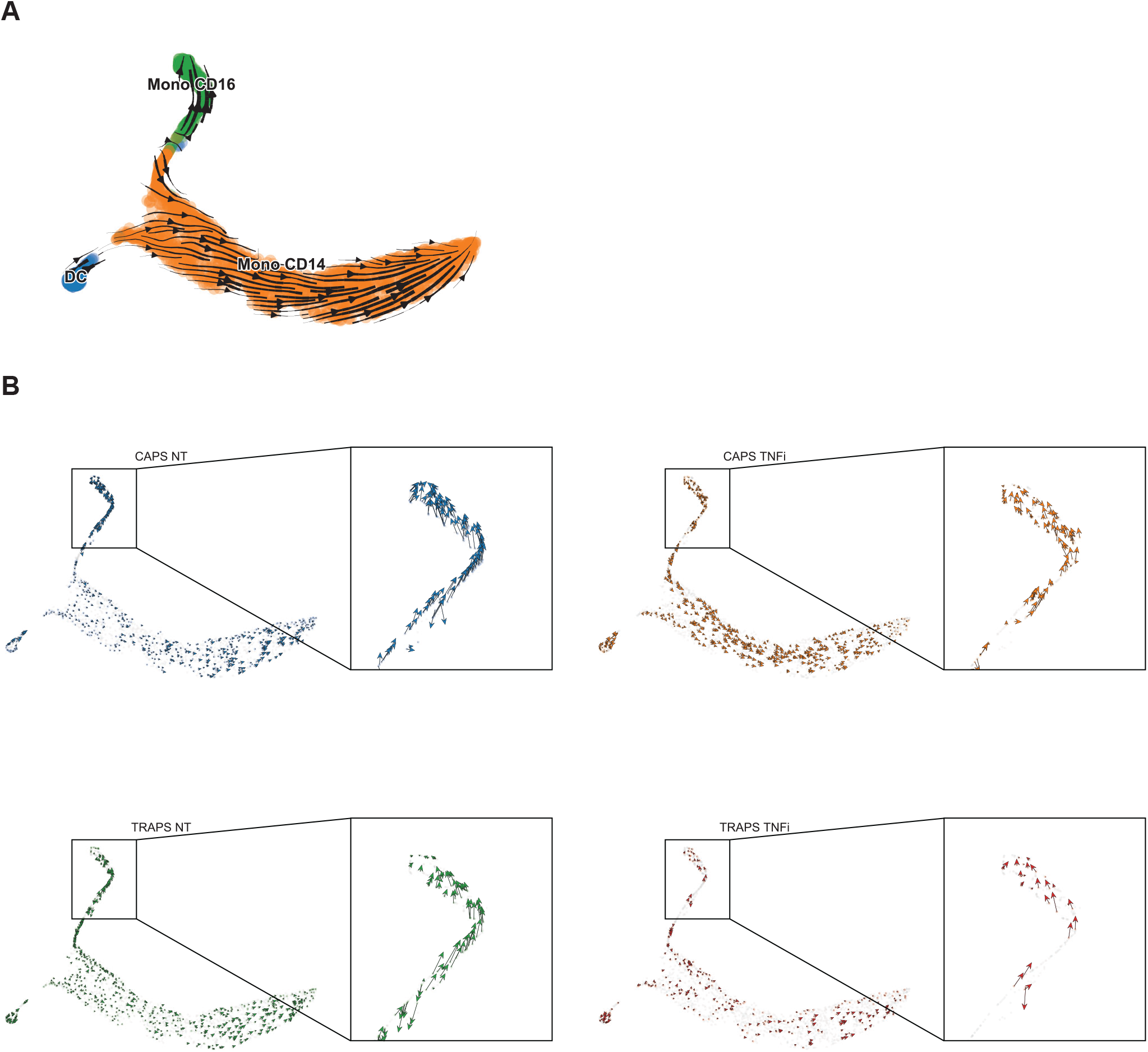

## 3 Discussion

In this study, we for the first time described the transcriptomic profile of PBMCs and PMNs in SAIDs patients, and also revealed transcriptional changes after anti-TNFα therapy. We observed that after anti-TNFα therapy, some pathways related to stress response and innate immune response-activating signal transduction were up-regulated, while some metabolic pathways related to aerobic respiration were down-regulated in CAPS, whereas there were no significant changes in TRAPS. This may suggest that the transcriptome is more associated with disease attack rather than disease activity, and may partially explain the unexpected good response to IL-1 blockade compared with anti-TNFα therapy observed in TRAPS patients in previous studies (Gattorno et al., 2008; Holzinger et al., 2015). We also identified that inflammatory cytokines such as TNFα and IL-1 β were up-regulated after anti-TNFα therapy and were not necessarily positively correlated with disease activity. Interestingly, some interferon-induced genes were up-regulated in the CAPS pre-treatment sample, which indicated that the interferon response can be correlated with disease flare. Further analysis revealed that one of the potential mechanisms of anti-TNFα treatment response in SAIDs may be the inhibition of macrophage differentiation rather than decreasing the overall level of inflammatory cytokines. As macrophages are a major cell type involved in inflammation, and their activation can be uncontrolled in SAIDs due to the genetic defect, inhibition of macrophage differentiation may help reduce the frequency of disease flare.

As SAIDs are rare diseases, in this exploratory study where we only enrolled 2 patients (one CAPS and one TRAPS) to test the feasibility of using scRNAseq for SAIDs research. As the number of samples was limited, we could not gain solid conclusions from this study. However, we can see that after anti-TNFα therapy, we observed significant transcriptional changes after treatment in CAPS patient, indicating that scRNAseq can be a useful technique for the future study of SAIDs and treatment response. Meanwhile, we are also aware of the shortcomings of this study design, which can be improved in the future: i) Some differentiated immune cells in the peripheral tissues (e.g. myeloid cells in skin lesions or synovial fluid) may have important pro-inflammatory functions. Therefore, we should consider not only extracting circulating PBMCs/PMNs but also collecting samples from peripheral tissues. ii) Some long-term effects caused by genetic variation or treatment may be the result of epigenetic modifications that cannot be revealed by scRNAseq, which is more reflective of immediate changes in cell state. Novel techniques such as single cell assay for transposase-accessible chromatin (ATAC) together with RNA sequencing may help us better understand such mechanisms. iii) In the study design we included neutrophils as they have important pro-inflammatory functions. However, we then realized that they had very few transcripts detected and less usable sequencing reads, as they had relatively low RNA content and relatively high levels of RNases with other inhibitory compounds. This brought great challenges to the subsequent analysis as we could not get high-quality information from these cells. In future studies, we should consider either supplementing RNase inhibitor to preserve the RNA or only keeping other high-quality cells without neutrophils.

In conclusion, we found that interferon response, rather than TNFα or IL-1, is involved in the disease flare of the SAIDs. Trajectory analysis showed that inhibition of macrophage differentiation as the potential mechanism of anti-TNFα treatment response in SAIDs. Our work provided new insights into the potential mechanisms of disease flare and anti-TNFα treatment response, new transcriptomic resources of SAID spectrum diseases, and expanded the application of single-cell RNA sequencing technology to new disease areas.

## 4 Materials and methods

### 4.1 Patients

This study enrolled 2 patients with classic SAIDs. One patient was a 20-year-old Chinese woman diagnosed as CAPS, with *NLRP3* D303G variation, and another patient was a 16-year-old Chinese man diagnosed as TRAPS, with *TNFRSF1A* V202D variation. Detailed information can be found in **Supplemental Data S1**. Both patients were in the cohort of adult SAID patients in Peking Union Medical College Hospital (PUMCH), which was described in the previous publication (Hua et al., 2019). This study was approved by the Institutional Review Board of Peking Union Medical College Hospital and performed according to the Declaration of Helsinki. Informed consents were obtained from all participants.

### 4.2 Single-cell RNA Sequencing

#### 4.2.1 Sample Preparation

The processing of blood samples, including PBMC and neutrophil isolation, library preparation, and single-cell RNA sequencing were done by CapitalBio Technology, Beijing. In brief, ACCUSPIN™ System-Histopaque^®^-1077 and Histopaque^®^-1119 reagent were used to isolate mononuclear cells and granulocytes, respectively, following the protocol on Sigma-Aldrich website. Using single cell 3’ Library and Gel Bead Kit V3 (10x Genomics, 1000075) and Chromium Single Cell B Chip Kit (10x Genomics, 1000074), the cell suspension (300-600 living cells per microliter determined by Count Star) was loaded onto the Chromium single cell controller (10x Genomics) to generate single-cell gel beads in the emulsion according to the manufacturer’s protocol. In short, single cells were suspended in PBS containing 0.04% BSA. About 6,000 cells were added to each channel, and the target cell will be recovered was estimated to be about 3,000 cells. Captured cells were lysed and the released RNA were barcoded through reverse transcription in individual GEMs. Reverse transcription was performed on a S1000TM Touch Thermal Cycler (Bio Rad) at 53°C for 45 min, followed by 85°C for 5 min, and hold at 4°C. The cDNA was generated and then amplified, and quality assessed using an Agilent 4200 (performed by CapitalBio Technology, Beijing).

#### 4.2.2 Data Processing and Cell Clustering

Raw sequencing data (fastq files) were mapped to the human genome (build GRCh38) using CellRanger software (10x Genomics, version 3.0.2). Raw gene expression matrices generated per sample were analyzed with the Seurat package in R (Stuart et al., 2019). To achieve clean cell clustering results, we divided the cell filtering process into two major steps: *primary clustering* and *fine adjustment. Primary clustering:* 5 samples (CAPS_NT, CAPS_TNFi, TRAPS_NT, TRAPS_TNFi, HD) were merged together and cells were filtered by nFeature_RNA (genes detected) > 200 and percent.mt (percentage of mitochondria genes) < 12.5%. High variable genes were selected by FindVariableFeatures and auto-scaled by ScaleData function using default parameters, and a principal component analysis (PCA) was performed for all datasets using the default RunPCA function in the Seurat package. Batch effect correction of each sample was done using the Harmony algorithm (Korsunsky et al., 2019) based on PCA space, followed by FindNeighbors and FindClusters function (dims = 30, resolution = 0.5) in the Seurat package for unsupervised clustering. In total 20 clusters were found. As plasma cells were not correctly identified by unsupervised clustering, they were manually annotated. *Fine adjustment*: scDblFinder package was used to predict potential doublets in the datasets (Germain et al., 2021). As neutrophils and platelets naturally have much fewer transcripts than other cell types, and DCs are often misclassified as doublets, we divided all cell types into 3 groups and use different filter criteria for each group. Group1: including Naïve CD4, Naïve CD8, Memory CD4, Memory CD8, Effector CD8, CD16^+^ NK, CD16^-^ NK, CD14 Mono, Naïve B, and Memory B, these cells were filtered by nFeature_RNA > 1200 and kept only singlets by scDblFinder; Group 2: including Neutrophil and Platelet, these cells were filtered by nFeature_RNA < 800 and kept only singlets by scDblFinder; Group 3, including FCGR3A Mono, DC, and pDC, these cells were filtered only by nFeature_RNA > 1200 regardless of scDblFinder prediction. In addition, 1 RBC cluster, 3 doublet clusters and 1 mitotic cluster were removed as they were not informative. All plasma cells were kept manually. After filtering, we ran the same pipeline as primary clustering mentioned above with slightly changes of several parameter (dims = 20, resolution = 0.4). Final clustering results were shown in Figure 1B.

#### 4.2.3 Differential Expression Analysis and Data Visualization

To visualize cells on a 2D plot, Uniform manifold approximation and projection (UMAP) was done by RunUMAP function (dims = 20) in Seurat. Differentially expressed genes (DEGs) were identified by the Wilcoxon Rank Sum test using the FindMarkers function. Gene expression levels or gene set enrichment scores (AUCell score) were shown in t-score or z-score for heatmaps or waterfall plots. UMAP or tSNE plots were done using DimPlot or FeaturePlot functions in Seurat. Heatmaps, modified stacked violin plots, boxplots, and bar plots of cluster proportion were generated using customized codes in R, and these functions were integrated into the R package “SeuratExtend” which is available on Github (https://github.com/huayc09/SeuratExtend).

#### 4.2.4 SCENIC and Gene Set Enrichment Analysis

To carry out the transcription factor network inference, SCENIC workflow was performed using Nextflow pipeline (Aibar et al., 2017), and regulon activity of each cell was evaluated using AUCell score with Bioconductor package AUCell. For functional/pathway analysis, gene set lists were collected from databases including Gene Ontology (GO) and Reactome. For Gene Set Enrichment Analysis (GSEA), the enrichment of given gene sets of each cell was evaluated using AUCell package as well. Gene regulatory networks (Figure S1B) were plotted using Cytoscape software (Shannon et al., 2003).

#### 4.2.5 Trajectory Analysis

For trajectory analysis (Figure 4), monocyte subsets of CAPS and TRAPS samples were extracted and integrated by Harmony algorithm. Python package Palantir (Setty et al., 2019) was used to calculate diffusion map and diffusion components based on Harmony space, then cells were visualized by tSNE. To predict the differentiation direction, we conducted a Velocyto pipeline (La Manno et al., 2018) using the *.bam file and barcode information generated by CellRanger, and used ScVelo in Python (Bergen et al., 2020) for better visualization.

### 4.3 ELISA

The plasma levels of IFN-α and IFN-γ were detected according to the instructions of the commercial ELISA kits (EXCELL Bio, China).

### 4.4 Statistical Analysis

Differentially expressed genes (DEGs) were identified by the Wilcoxon Rank Sum test using the FindMarkers function of Seurat package in R. All remaining statistical analysis were performed by the ggpubr package in R with default parameters.

## 6 Author Contributions

M.S. and Y.H. conceived the study. Y.H. performed the bioinformatics analyses. N.W. performed the Elisa experiment. J.M. summarized the medical history of SAID cohort. Y.H. and M.S. wrote the manuscript. All authors discussed the results and commented on the manuscript.

## 8 Acknowledgments

The authors thank CapitalBio Technology for the technical support of single-cell RNA sequencing; Peking Union Medical College Hospital Laboratory Department for testing serum cytokines; Dr. Junbin Qian (Zhejiang University) for the suggestions of single cell analysis; Dr. Kathryn Jacobs (VIB-KU Leuven) for the help with language modification.

## 9 Supplementary Material

Supplemental Figure S1. Interferons and Interferon Signaling Pathways

Supplemental Data S1. Case presentation

## 10 Data Availability Statement

The raw sequence data reported in this paper will been deposited in the Genome Sequence Archive (Chen et al., 2021) in National Genomics Data Center (CNCB-NGDC Members and Partners, 2022), China National Center for Bioinformation / Beijing Institute of Genomics, Chinese Academy of Sciences that are publicly accessible at https://ngdc.cncb.ac.cn/gsa. All software is freely or commercially available.

