## Supplemental Figure S1 for "Single-cell Transcriptomic Analysis of Systemic Autoinflammatory Diseases with Anti-TNFα Therapy"

### Supplemental figure 1

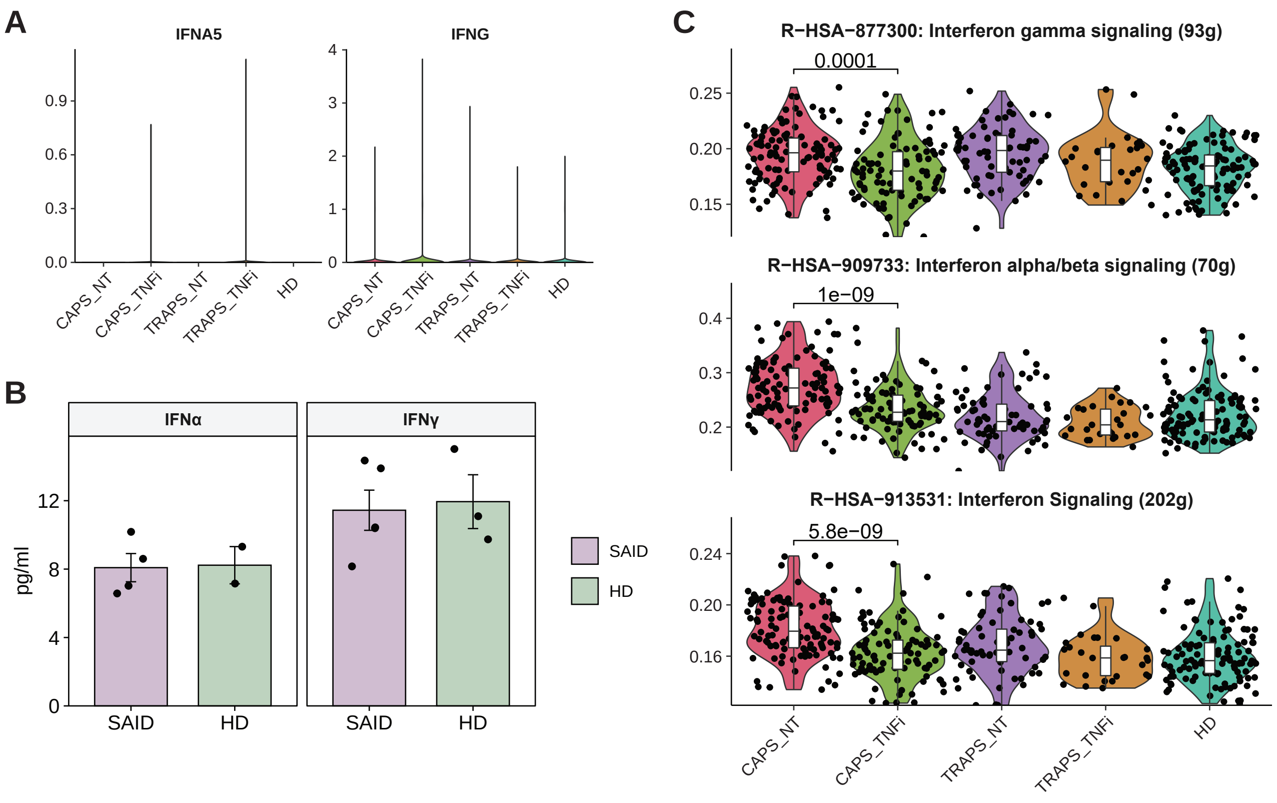


**Figure S1. Interferons and Interferon Signaling Pathways.**

(A) Violin plots showoing the expression of IFNs in all cells, comparing 5 sample origins.

(B) Serum IFNα and IFNγ levels in non-treated SAID patients and healthy donors by ELISA.

(C) Violin plots showing the AUCell enrichment scores of interferon signaling gene lists from Reactome database in CD16+ monocytes, comparing 5 sample origins.
