## Supplemental Data S1 for "Single-cell Transcriptomic Analysis of Systemic Autoinflammatory Diseases with Anti-TNFα Therapy"

**Supplemental Data S1. Case presentation**

Patient 1

A 20-year-old Chinese woman reported persistent cold-triggered fever, polyarthritis and urticaria-like rash after birth, usually lasting for 1h and relieving spontaneously. She also suffered from papilledema, vision impairment and sensorineural hearing loss two years ago. High serum ESR (70 mm/h), CRP (112 mg/L) and IL-6 (6.5 pg/ml) levels were observed in the attack phases. Her mother had similar symptoms of rashes, arthritis, ophthalmia and became blindness eventually. A heterozygous *NLRP3* (NM_001243133.1) gene mutation c.908A>G (p.D303G) was confirmed in the proband and her mother. She was diagnosed with CAPS, and treated with TNFα inhibitor (etanercept) plus methotrexate, with a satisfactory response to fever, polyarthritis, dermatitis and impaired vision. ESR, CRP and IL-6 levels also decreased after therapy.

Patient 2

A 16-year-old Chinese man suffered from recurrent fever, rashes and abdominal pain since the age of one, accompanied by headache, arthralgia, myalgia, conjunctivitis and periorbital edema. The disease attacked every several weeks, with each attack lasting 2-3 days. The acute phase reactants elevated (ESR 68 mm/h and CRP 200 mg/l) during the flares while decreased (ESR 30mm/h and CRP 48.3mg/L) during the intervals, yet not completely to the normal level. Genetic testing identified a de novo heterozygous variant in the *TNFRSF1A* gene (NM_001065), c.605T>A (p.V202D). He was diagnosed with TRAPS and administrated with etanercept. His symptoms of fever, rashes, arthralgia and abdominal pain relieved at subsequent follow-ups.
